# Renewed diversification following Miocene landscape turnover in a Neotropical butterfly radiation

**DOI:** 10.1101/148189

**Authors:** Nicolas Chazot, Keith R. Willmott, Gerardo Lamas, André V. L. Freitas, Florence Piron-Prunier, Carlos F. Arias, Jim Mallet, Donna Lisa De-Silva, Marianne Elias

## Abstract

“This preprint has been reviewed and recommended by Peer Community In Evolutionary Biology (http://dx.doi.org/10.24072/pci.evolbiol.100032)”

The Neotropical region has experienced a dynamic landscape evolution throughout the Miocene, with the large wetland Pebas occupying western Amazonia until 11-8 my ago and continuous uplift of the Andes mountains along the western edge of South America. Although the complex dynamics between the Andes and Amazonia may have strongly affected the trajectory of Neotropical biodiversity, there is little evidence for such an influence from time-calibrated phylogenies of groups that diversified during this period. Here, we generate one of the most comprehensive time-calibrated molecular phylogenies of a group of Neotropical insects: the butterfly tribe Ithomiini. Our tree includes 340 species (87% of extant species), spanning 26 million years of diversification in the Neotropics. We investigate temporal and spatial patterns of diversification, focusing on the influence of Miocene landscape tranformations on the dynamics of speciation, extinction and biotic interchanges at the Amazonia/Andes interface. We find that Ithomiini likely began diversifying at the interface between the Andes and the Amazonia around 26.4 my ago. Five subtribes with a very low extant diversity started diversifying early in western Amazonia, but a rapid decrease in diversification rate due to increased extinction rate between 20 and 10 my ago suggests a negative impact of the Pebas wetland system on these early lineages. By contrast, the clade containing the five most species-rich subtribes (85% of extant species) was characterized by a high, positive net diversification rate. This clade diversified exclusively in the Central Andes from 20 to 10 my ago. After the demise of the Pebas system (11-8 my ago), we found a sudden increase of interchanges with the Northern Andes and Amazonia, followed by local diversification, which led to a substantial renewal of diversification. In general, ecological turnovers throughout the Miocene strongly determined the dynamics of speciation, and extinction and interchanges, and appear as a key driving force shaping the region’s current extraordinary biodiversity.

## INTRODUCTION

There has been a long fascination among biologists for the Neotropics and the origin of its intriguingly high biodiversity. The timing of Neotropical diversification, and therefore its major driving processes, is still controversial despite the large amount of publications that have addressed the question (e.g., ^1,2,3^)

Despite uncertainty about the precise timing and magnitude of surface uplift, the formation of the Andean cordilleras during the Cenozoic greatly shaped Neotropical landscapes and affected diversification in the Neotropics. As the Andes arose, they brought about new biotic and abiotic conditions along their slopes, modified the climate of the Neotropical region and deeply affected the formation of the Amazonian basin by bringing large amounts of sediments and modifying water drainage^1^. There is increasing evidence that the Andes influenced the diversification of Neotropical lineages, primarily by increasing speciation rate, perhaps most spectacularly in the high altitude páramo habitat (e.g., ^4^). In parallel, the western part of the Amazon basin, which is connected to the Andes, has experienced major turnovers of ecological conditions. During the Oligocene, western Amazonia was occupied by a fluvial system flowing northward (paleo-Orinoco basin), which transformed ~23 million years (my) ago into an aquatic system of shallow lakes and swamps episodically invaded by marine conditions, known as the Pebas system ^1,5,6,7,8^. The Pebas was connected northward with the Caribbean Sea and likely also with the Pacific Ocean through the Western-Andean portal (“WAP”, ^5^), a low-altitude gap that separated the Central Andes and the Northern Andes until 13-11 my ago. During the late Miocene and during the Andean uplift, the accumulation of sediments combined with a sea level decrease initiated the eastward drainage of the Pebas, and by 10-8 my ago the region had changed into a fluvial system, which then turned into the modern configuration of the Amazon. More recently, climatic fluctuations during the Peistocene (2.5-0 my ago) may have led to episodic dryness affecting Amazonian forest habitats ^9^. The extent of the influence of Pleistocene events and their effects on Neotropical diversification, and even the importance of dryness episodes, are controversial (e.g,^2,10,11,12^)

In this study we purposely focus our attention mostly on the Miocene and Pliocene and how the interaction between the rise of the Andes and coincident large landscape modifications in western Amazonia have determined diversification and dispersal over 30 million years. The Pebas ecosystem covered up 1.1 million km^2^ at its maximum ^7^ and was probably not suitable for a terrestrial fauna. Therefore, between 23 and 10 my ago, diversification of terrestrial lineages may have been impeded in western Amazonia or restricted to its edges ^7^. By contrast, the uplift of the Central and the Northern Andes, also occurring throughout the Miocene and the Pliocene, and the ecological gradients present along this mountain chain probably constituted an important driver of diversification. In the last 10 to 8 my, the retreat of the Pebas may have provided opportunities for terrestrial lineages to radiate in western Amazonia. The Pebas may also have constrained dispersal, acting as a barrier between the Andes, the Guiana shield and western Amazonia. Thus, determining whether rates of interchanges have been constant throughout time since the origin of Ithomiini, or instead have increased after the Pebas’s retreat, would test the potential importance of this ecosystem in building the modern pattern of diversity ^6^.

Paleontological studies have shown that the Pebas greatly contributed to the diversification of aquatic fauna such as molluscs ^13^, ostracods ^14^ and crocodilians ^15^. However, the fossil record also suggests a negative effect of the Pebas system on terrestrial fauna ^16,17^. The hypothesis that the Pebas has shaped patterns of terrestrial diversification and dispersal in western Amazonia has grown over the years (e.g., ^5,6,7,16,18^) but support from molecular phylogenies mostly stems from the observation that western Amazonian clades have diversified during the last 10-8 my and not before (^19,6^ and reference therein). Yet, there is very little information on what happened before, when the Pebas was occupying western Amazonia, particularly on whether the presence of Pebas constrained diversification and interchange patterns in this region. A thorough assessment of the role of the Pebas ecosystem on diversification and dispersal requires phylogenies of large Neotropical clades that originated before the formation of the Pebas, i.e. clades older than 23 my old. Phylogenies of Neotropical clades meeting these conditions are surprisingly rare. In insects – which are among the most diverse terrestrial organisms - attempts to build phylogenies of Neotropical groups to test different drivers of Neotropical diversification have either suffered from a small size or a low sampling fraction (e.g.^19,20,21,22,23^) and therefore from low statistical power and reliability.

Butterflies are among the best candidates for addressing the evolution of the Neotropical biota at such time scales. They are probably the best-known insects and Neotropical butterfly lineages have benefited from substantial phylogenetic research compared to other insects (e. g., ^19,20,21,24,25,26,27,28,29,30,31,32,33,34,35^). Among the most emblematic Neotropical butterflies is the tribe Ithomiini (Nymphalidae: Danainae, 393 species), also referred to as the clearwing butterflies because of the transparent wings of the majority of species. Ithomiini are forest-dwellers distributed throughout the Neotropics, from sea level up into montane cloud forests (to 3000 m), where their larvae feed on plants of the families Solanaceae, Gesneriaceae and Apocynaceae ^36^. Species richness is primarily concentrated in the Andes, where about half of the species occur (mostly on the eastern slopes) and in western Amazonia. Ithomiini are chemically defended and they engage in Müllerian mimicry, whereby co-occurring species exhibit convergent wing colour patterns that advertise their toxicity to predators ^37^. Ithomiini butterflies represent a keystone group in Neotropical forests by numerically dominating mimetic butterfly communities and sharing wing colour patterns with a large number of other palatable and unpalatable Lepidoptera, such as the iconic *Heliconius* butterflies ^38^. For this reason, Ithomiini were used by both Bates ^39^ and Müller ^37^ in their original descriptions of deceptive (Batesian) and mutualistic (Müllerian) mimicry, respectively.

The diversity and the intriguing ecology of Ithomiini has generated a great interest and a broad and diverse literature on topics including life history ^40,41,42,43^, chemical ecology ^44,45,46^, systematics ^19,21,26,32,34,36,47,48^, cytogenetics ^49^, community ecology ^50,51,52,53,54,55^, wing colour pattern evolution ^56^, and biogeography ^19,21,26,27^. In this study, we build on existing molecular data and provide a large amount of novel DNA sequences for ithomiine species to generate the first species-level phylogeny of the entire tribe, providing a large and densely sampled (340 species included out of 393 currently recognized) phylogenetic dataset for a Neotropical insect clade that underwent diversification during the last ~30 million years ^27,57^. This phylogenetic framework provides an ideal opportunity for investigating Neotropical diversification over a large evolutionary time-scale. Ithomiini originated before the Pebas system, thus offering the opportunity to investigate diversification before, during and after the environmental changes during the Miocene with a high statistical power. Importantly, contrary to many other large radiations of similar age, Ithomiini are endemic to the Neotropics. Their diversification therefore occurred without interaction with other biogeographic regions such as the Nearctic.

Here, we investigated the dynamics of speciation, extinction and dispersal rates in Ithomiini through space and time, using a combination of time- and trait-dependent models of diversification and historical biogeography. We focused on the interaction between the turnover of ecological conditions in western Amazonia and the Andean uplift during the Miocene, and we investigated whether geological and ecological events in both regions affected synergistically the diversification of Ithomiini. More specifically, an important role for Andean uplift and the Pebas would be supported if: (1) *During the Pebas period*: (a) Andean diversification largely exceeds Amazonian diversification, due to an increased diversification in the Andes driven by the evolving ecological gradient and uplift dynamics and/or a reduced diversification rate in Amazonia accompanying the loss of terrestrial habitats; (b) interchanges with western Amazonia are reduced; (c) interchanges between the Central and the Northern Andes are reduced, because of the existence of the WAP. (2) *During the retreat of the Pebas*: interchanges with western Amazonia and between the Central and the Northern Andes largely increase, as a result of new terrestrial habitats and the disappearance of the WAP, respectively. (3) *After the Pebas period*: Diversification rates in Amazonia globally increase and biotic interchanges are not constrained anymore. After the Pebas retreat, decrease of speciation rates through time suggest post-Pebas radiations in Amazonia, while increase of speciation rates through time may suggest a role of climatic fluctuations during the last 2.5 my.

## RESULTS

### Time-calibrated phylogeny

#### Tree topology and time-calibration

We generated a time-calibrated phylogeny that comprised 340 out of 393 Ithomiini species (Supporting Information S1-S2-S3), based on three mitochondrial and six nuclear gene fragments. The tree topology was obtained under maximum likelihood inference, and branch lengths were estimated by Bayesian inference, using six secondary calibration points from Wahlberg et al. (^57^). The tree topology was generally well supported, including deep nodes (Supporting Information S4). We found a crown age of Ithomiini of 26.4 (CI=22.75-30.99) my ago (Figure 1, Supporting Information S5) and a divergence time from its sister clade Tellervini of 42.1 (CI=39.50-48.44) my ago. All subtribes (10 in total) diverged in the first 10 million years, in the following order (Figure 1): (1) Melinaeina (26.4, CI=22.75-30.99 my ago), (2) Mechanitina (24.2, CI=21.00-28.59 my ago), (3) the clade consisting of Tithoreina and the Methonina (23.6, CI=20.40-27.71 my ago), (4) Athesitina (22.1, CI=19.09-26.25 my ago) and (5) a large clade that comprises the five most species-rich subtribes (Ithomiina, Napeogenina, Oleriina, Dircennina and Godyridina), hereafter called the “core-group”. The relationships between tribes were similar to those recovered in a recent higher-level phylogeny of Ithomiini based on a combination of 3 gene regions and morphological characters ^34^, except that Brower *et al.* (^34^) recovered Mechanitina as sister to Tithoreina+Methonina. Lineage ages in our phylogeny were generally younger than those inferred in Wahlberg *et al.* (^57^), but older than those inferred in Garzon-Orduña *et al.* (^58^) (see De-Silva *et al.* ^48^ for further discussion of such differences).

**Figure 1.**
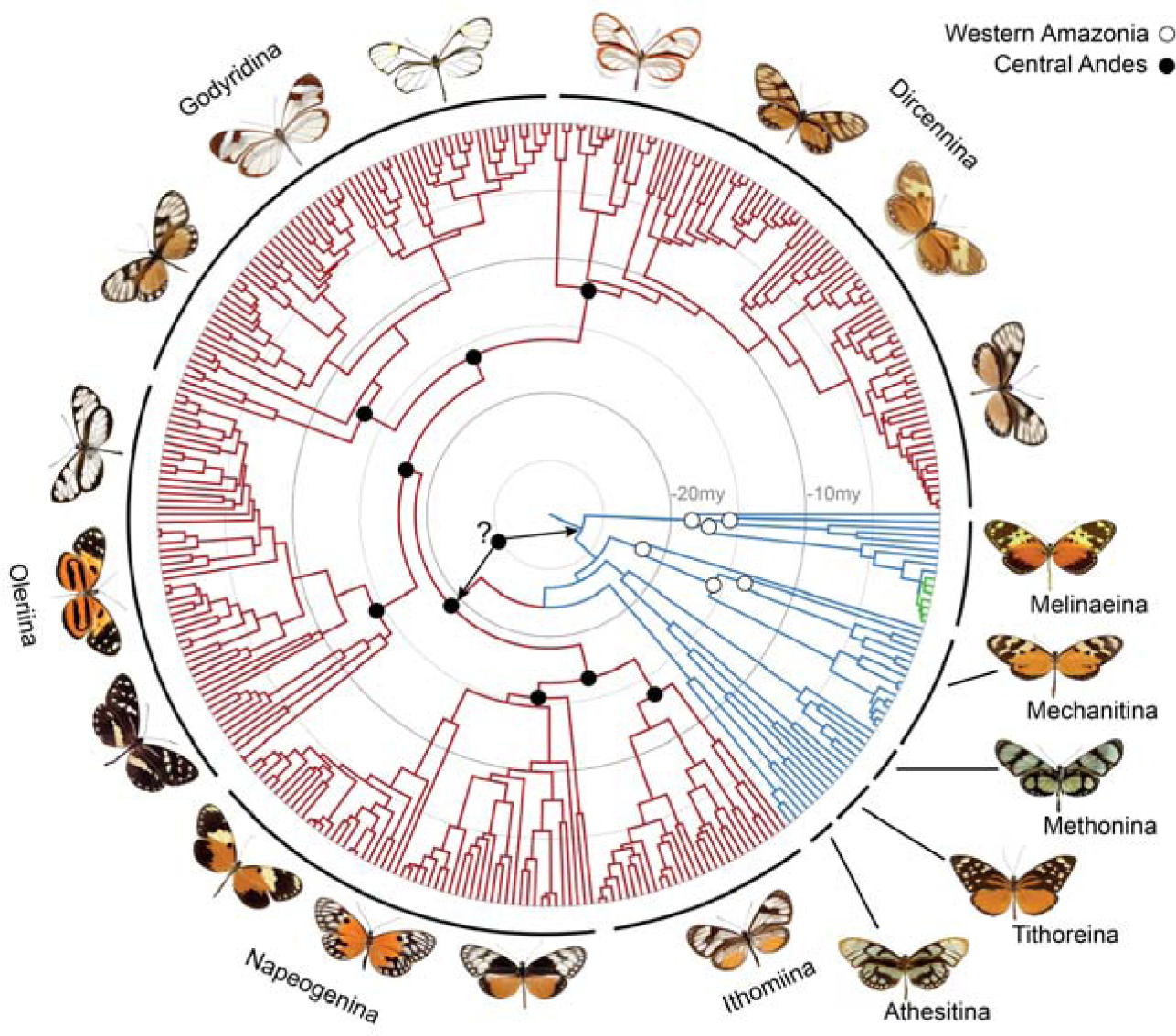
Time-calibrated phylogeny of the Ithomiini tribe. Coloured branches depict the partitions identified by MEDUSA and used for fitting diversification rate models. Red lineages constitute the core-group, green lineages the *Melinaea*-group, blue lineages the background tree. Black and white circles indicate the biogeographic ancestral states reconstructed at the basal nodes of the tree: black=Central Andes, white=Western Amazonia. Question mark and arrows indicate the position of two alternative scenarios for the first colonization of the Andes: BioGeoBEARS at the root of the Ithomiini, BiSSE at the root of the core-group. Both methods agree for Melinaeina and Mechanitina but diverge for Tithoreina, Methonina and Athesitina. The names and position of the different subtribes are indicated.

### Diversification rates

#### Time-dependent diversification

We combined three methods to assess patterns of diversification through time and across clades. We first used MEDUSA ^59^, which automatically detects shifts of diversification processes across a phylogeny. The analysis detected two significant shifts from the background process of diversification (Figure 1, Supporting Information S6). One was at the root of the core-group accounting for ~85% of present-day diversity of Ithomiini. The second shift was at the root of a subgroup within the genus *Melinaea,* which appears to have diversified rapidly during the last one million years (hereafter “*Melinaea*-group”).

We then used the method developed by Morlon *et al.* ^60^ to fit time-dependent models of speciation and extinction on the different partitions based on the MEDUSA results. This method does not detect shifts automatically, but it allows both speciation and extinction rates to vary through time and across lineages. The results confirmed that the partitioned models (either or both of the two shifts detected by MEDUSA) had a significantly better fit and that the two-shift model was significantly better than the one-shift models (Table 1). For the core-group, no model of time-dependent diversification had a significantly better fit than the null model of constant speciation rate without extinction (0.255 lineage^-1^ my^-1^, Figure 2, Table 1, Supporting Information S6). Under the null model, the core-group diversity increased exponentially during the last 20 million years and reached its current diversity (334 extant species). For the *Melinaea*-group, the best fitting model was an exponentially decreasing speciation rate without extinction, with a very high initial speciation rate (7.62 lineage^-1^my^-1^ at the root, 0.342 lineage^-1^my^-1^ at present, Figure 2, Table 1, Supporting Information S6). The *Melinaea*-group radiated into eight species during a time lapse of only one million years. On the remaining background lineages, the best fitting model involved a time-dependency of both speciation and extinction rates. The resulting diversification rate was high during the initial stages of diversification (0.75 at the root), but decreased rapidly and became negative around 19 my ago. The background diversification rate then started a slow recovery, but remained negative (-0.038 lineage^-1^my^-1^ at present, Figure 2, Table 1, Supporting Information S6). Consequently the background tree diversity reconstructed from this model shows a pattern of diversity that increased during the first 10 my up to ~60 species before slowly declining toward its current diversity (51 species) during the last 15 my. This signal of high relative extinction rate was not affected by initial parameters for maximum likelihood search or by diversification rate heterogeneity potentially remaining from the background (see discussion in Supporting Information S7). Clades within the background tree showing positive net diversification rates during the last 5 my supported the recovery trend described above (Supporting Information S7).

**Table 1.**
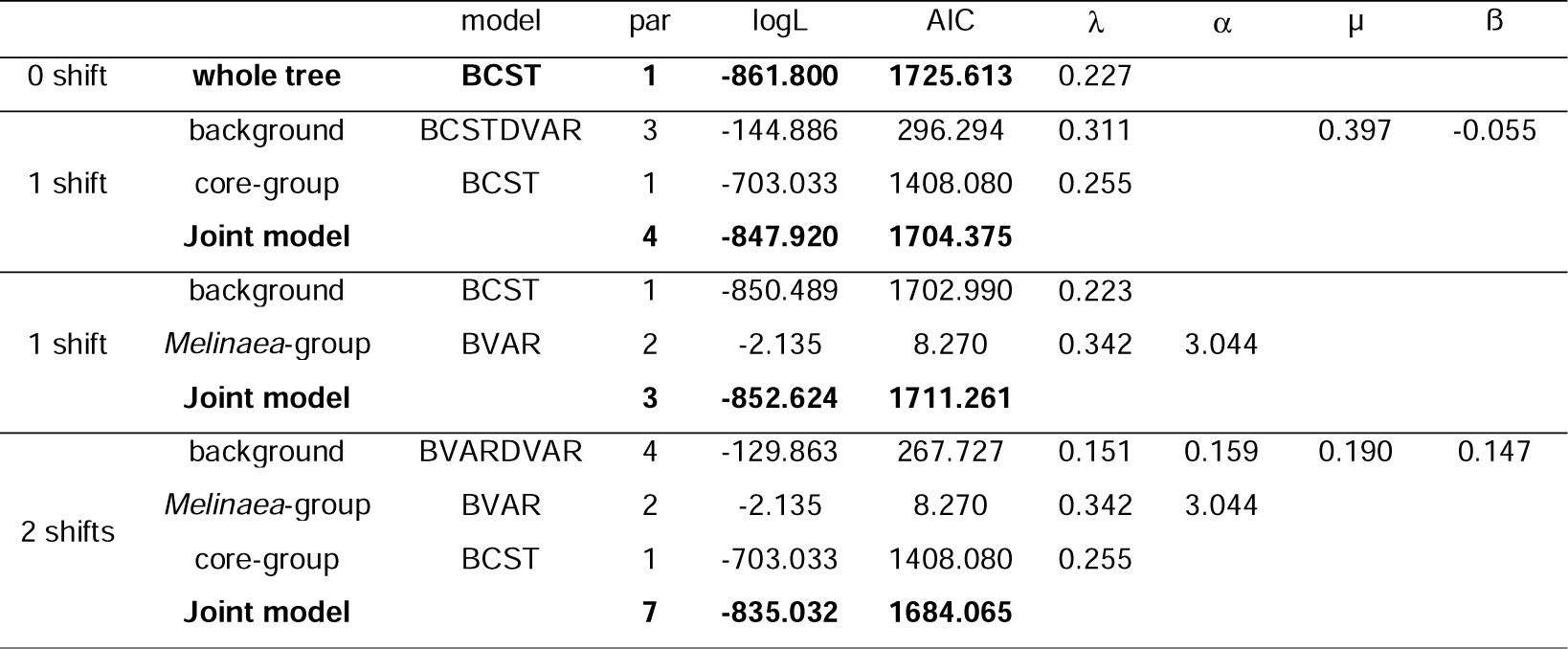
Results of time-dependent models of diversification fitted on the different partitions: 0 shift, 1 shift (core-group or *Melinaea*-group), 2 shifts. For each subclade or the background tree, only the best fitting model is shown (see Supporting Information S6 for more details). BCST=constant speciation, BVAR=time-dependent speciation, DCST=constant extinction, DVAR time-dependent extinction. λ =speciation rate at present, α=coefficient of time variation of the speciation rate, μ =extinction rate at present, ß =coefficient of time variation of the extinction rate.

**Figure 2.**
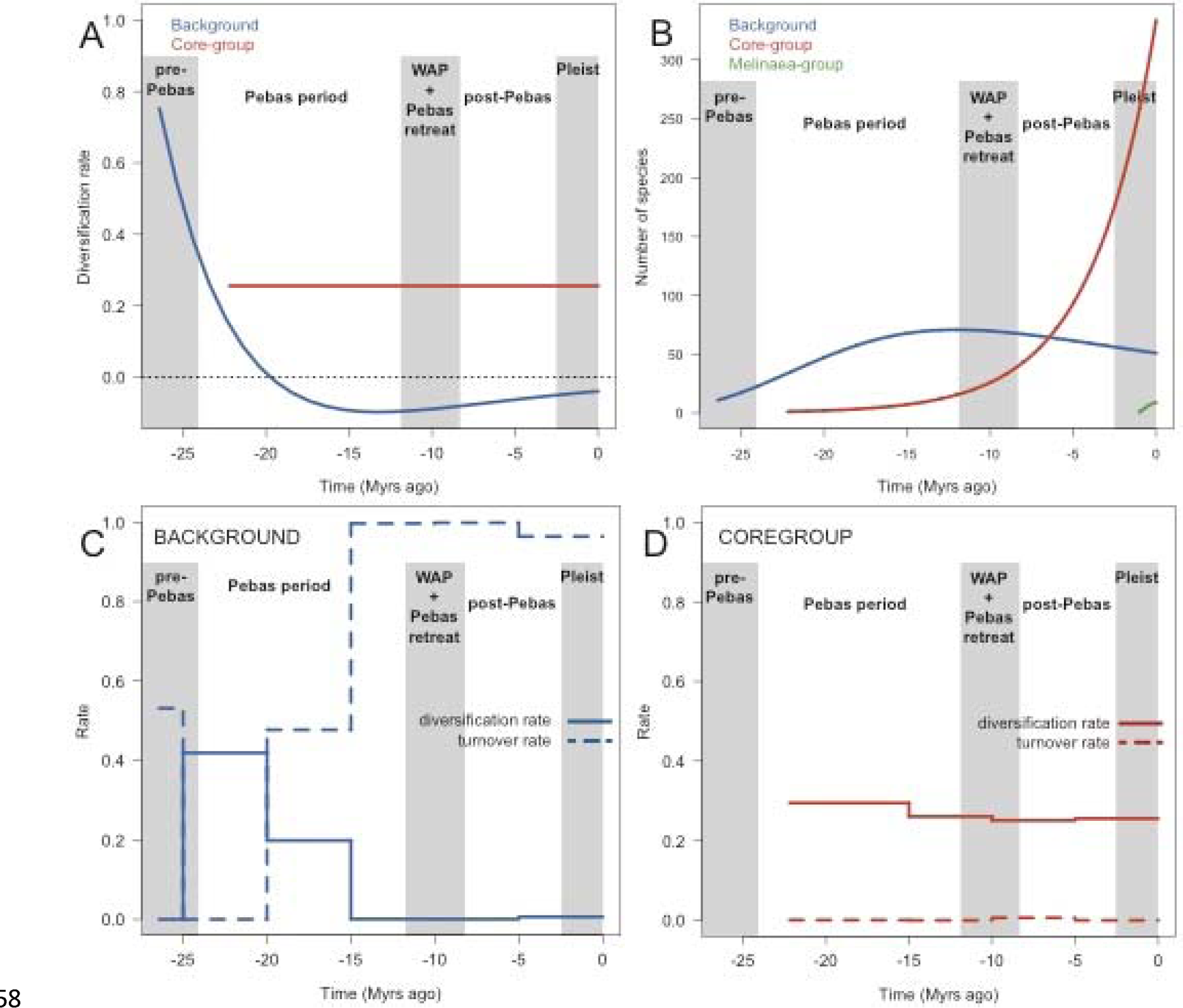
Results of diversification rate estimates through time using MEDUSA’s partition. A. Diversification rate estimated using the method designed by Morlon *et al.* (^60^) for the background and the core-group. Results for the *Melinaea*-group are not plotted because of its very high diversification rate. B. Diversity trajectories inferred from the best fitting models of diversification obtained using Morlon *et al.* (^60^)’s method. C. and D. Diversification rate and turnover of the background and coregroup respectively, estimated using 5 million-year time-bins with TreePar ^61^.

We also used TreePar ^61^ to fit models of constant diversification rate in 5 my time-bins for each partition identified by MEDUSA, as a second and independent assessment of diversification through time. TreePar accommodates constant birth-death models within time-intervals and allows those birth-death rates to vary between time intervals. The results were congruent with those of the time-dependent models of diversification obtained with Morlon *et al.* (^60^)’s method (Figure 2, Supporting Information S7). The diversification rates estimated for the core-group within 5 my time-bins remained relatively constant through time and the turnover rate was close to 0, supporting a null or extremely low relative extinction. For the background tree, we found that diversification rate was highest between 25 and 20 my ago, declining toward 0 during the last 15 my, in agreement with the results obtained with the method of Morlon *et al.* (^60^). Turnover largely increased and reached a maximum during the last 15 million years, supporting a very high relative extinction rate, similarly to our results of the time-dependent diversification models of Morlon *et al.* (^60^)’s method.

We additionally performed an analysis with BAMM v.2.5.0 ^62^, which also detects shifts in diversification dynamics using a Bayesian framework. The results supported the presence of a shift at the root of the core-group with an increasing diversification rate compared to the lineages diverging earlier. More details about this analysis can be found in Supporting Information S8.

#### Diversification in the Andes

We compared the pattern of diversification in the Andes and in the rest of the Neotropics and we assessed the rate of interchanges between these regions using character state-dependent diversification models (ClaSSE, ^63^). Such models estimate character state-dependent rates of speciation, extinction and cladogenetic state transitions (i. e., occurring at nodes). Species were classified into two biogeographical states, Andean and non-Andean. We compared 10 models to test whether rates of speciation, extinction or transitions were different or not between regions. For both the whole Ithomiini and the core-group we failed to identify a single model that fitted the data significantly better than other models (Table 2). For the whole Ithomiini, the model with the lowest AIC score had a higher speciation rate in the Andes (λ_222_=0.230) than in non-Andean regions (λ_111_=0.118) and those speciation rates were higher than the colonization rates (λ_112_=λ_212_=0.079). The second best model had, again, a higher speciation rate in the Andes (λ_222_=0.231) than in non-Andean regions (λ_111_=0.108) and the colonization rate out of the Andes was higher (λ_212_=0.095) than into the Andes (λ_212_=0.047). In the core-group, there were four models within an AIC difference of 2. All models had extremely low extinction rates. Only one of them inferred different speciation rates and three of them inferred a higher colonization rate into the Andes than out of the Andes (Table 2). Therefore, there is no support for a general pattern of increasing diversification in the Andes across the entire core-group (˜85% of the extant diversity). This interpretation was confirmed by additional analyses in which we tested the presence of a hidden character using the model HiSSE ^64^ The results supported the conclusion that the Andes are not directly associated with higher diversification rates (see Supporting Information S9), and that diversification was also fast in some non-Andean clades and slow in some Andean lineages.

**Table 2.**
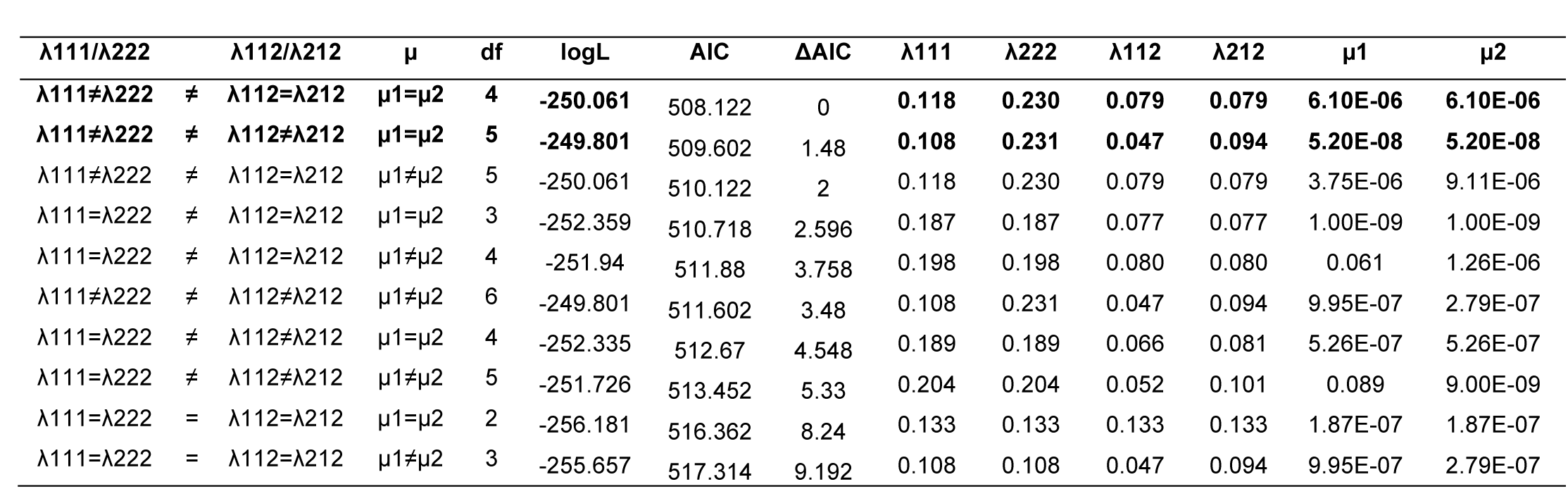
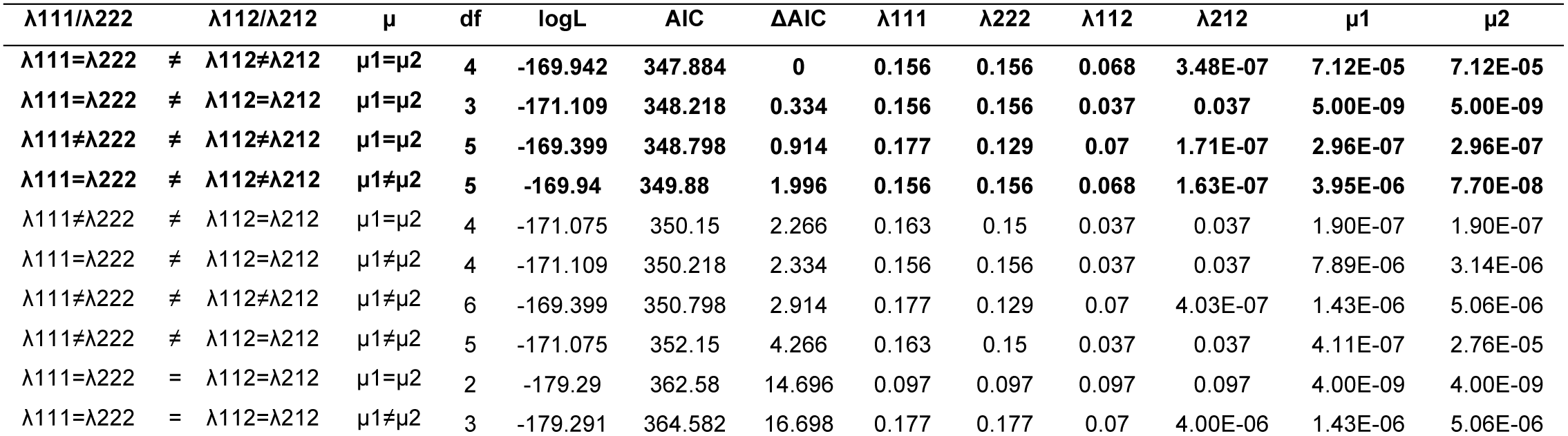
Results of ClaSSE models fitted on the whole Ithomiini tree (a.) and the core-group (b.) sorted by increasing AIC. Constraints of each model are indicated in the four first columns. 1=non-Andean, 2=Andean, λ111 and λ222 represent within-region speciation rates, λ112 and λ212 represent cladogenetic transition rates, μ = extinction rates, df = degree of freedom (number of parameters), logL=log-likelihood, AIC= Akaike information criterion score, ΔAIC = AIC difference with the best fitting model.
a. Whole tree
b. Core-group

Ancestral state inference is not implemented in ClaSSE. Instead, we used the BiSSE ^65^ model, in which transitions occur only along branches. To infer ancestral states on the whole tree, we fitted the BiSSE models corresponding to the best-fitting ClaSSE models (i.e. model1: different speciation rates, model2: different speciation and transition rates). In both cases the most likely state of the root was non-Andean but with high uncertainty (probability of 0.508 and 0.543 respectively) and there was uncertainty at the nodes leading to the core-group (Figure 1, Supporting Information S10). The most recent common ancestor (hereafter MRCA) of the core-group was inferred as most likely Andean (model1: probability of 0.558, model2: probability of 0.625). In both models the MRCA of all background subtribes were inferred to be non-Andean with a strong support (except for Athesitina, Figure 1, Supporting Information S10). In the core-group, the best model (with different speciation rates) inferred an Andean origin for subtribes Napeogenina, Dircennina and Godyridina, whereas the second best model (different speciation and transition rates) inferred an Andean origin for all five subtribes (Supporting Information S10).

#### Diversification in Amazonia

We also investigated the pattem of Amazonian diversification during the post-Pebas period (8-0 my ago). We fitted a model of time-dependent speciation rate using Morlon et al.’s method ^60^ (no extinction, based on our previous results) to assess whether speciation rates decreased through time (supporting radiations accompanying the post-Pebas recolonizations) or increased through time (supporting a recent diversification potentially caused by Pleistocene climatic fluctuations). We identified four Amazonian diversification events – clades whose nodes were inferred to be almost all Amazonian (see historical biogeography results below) – in the core-group and three in the background subtribes. All core-group clades showed decreasing speciation rates through time, suggesting an early diversification, perhaps following the appearance of new forest habitats accompanying the Pebas retreat. Among the three clades from the background lineages only the genus *Methona* followed a trend of decreasing speciation rate. The other two clades, *Mechanitis* + *Forbestra* and *Melinaea* (the whole genus) supported an increasing speciation rate through time, which is consistent with a potential effect of Pleistocene climatic fluctuations in driving diversification.

### Historical biogeography

We divided the Neotropical region into 9 areas (Supporting Information S11) and we assigned each ithomiine species to these areas according to their current distribution. We performed historical biogeographic reconstruction using BioGeoBEARS ^66^, under two models (DEC and DIVALIKE), using a three-step procedure outlined below. We first performed a “null” model with uniform dispersal multipliers. Based on the results of the null model, we computed rates of dispersal between specific regions for all 1-my intervals. We then implemented those rates in a time-stratified model.

#### Biogeographic null model

We started with a “null” biogeographic model, which restricted the area adjacency but set all dispersal probabilities to 1, and we compared the models DEC and DIVALIKE. The null DEC model had a better fit than the null DIVALIKE model (likelihoods: DECnull: −1335.802, DIVALIKEnull: −1347.869), hence we used the DEC model in all subsequent analyses.

In both models, the ancestral area of the Ithomiini MRCA was unclear (Central Andes + Western Amazonia for the highest probability). The areas where the two first divergences occurred, which led to the Melinaeina and the Mechanitina subtribes, were also unclear (Figure1, Supporting Information S12). The ancestor of the remaining ithomiine lineages was recovered to be only occupying the Central Andes. Following this node (23.6 my ago) all the divergences occurred in the Central Andes until 9.4 my ago, when the first colonization event out of the Central Andes occurred (MRCA of *Oleria*, which dispersed into Western Amazonia). Hence, our null biogeographic reconstruction found that all subtribes except Melinaeina and Mechanitina originated and started diversifying in the Central Andes. Interchanges between regions appeared to have increased during the last 10 my. However, all node reconstructions at the basal nodes of the background lineages were highly uncertain.

#### Biogeographic diversification of the core-group

Using this null model, we investigated more specifically the biogeographic pattern of the core-group by computing rates of dispersal among different regions. We applied the null model to 100 trees randomly sampled from the BEAST posterior distribution and extracted the state with the highest probability at each node. Then, for each 1-my interval, we computed the number of specific transitions divided by the number of lineages existing during this interval and fitted a spline line on the distribution of points.

As observed in the ancestral state reconstruction on the maximum clade credibility tree (MCC), no dispersal event occurred during the initial Central-Andean phase of diversification in the core-group (Figure 3). Between 13-8 my ago a major peak of interchanges between the Andes and Amazonia occurred, followed by a second peak between 4-0 my ago (Figure 3). The first peak was almost entirely driven by colonization from the Andes toward the Amazonia, whereas the second peak involved many reverse colonizations toward the Andes. We also recorded a large peak of colonizations from the Central Andes toward the Northern Andes between 13-8 my ago, also followed by a second peak 4-0 my ago (Figure 3, Supporting Information S11-S12). Colonization of Central America may have started 8 my ago, but interchanges largely increased during the last 4 my (Supporting Information S11). Colonizations of the Atlantic Forest also started early (around 13 my ago), but the rate of interchanges between the Atlantic Forest and the remaining Neotropical regions remained relatively constant during the last 10 my (Supporting Information S11-S12).

**Figure 3.**
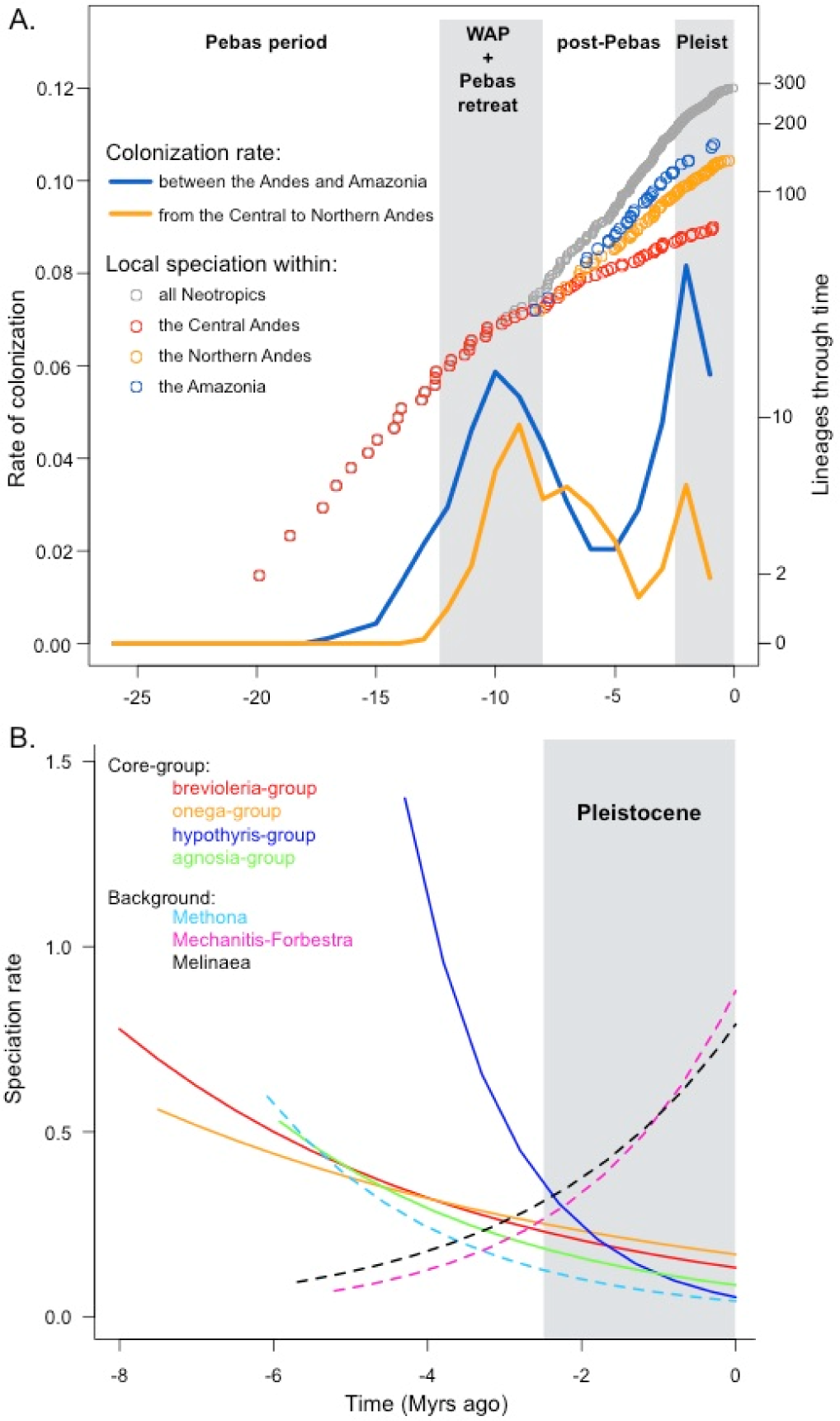
A. Colonization rates and lineage accumulation through time (by speciation) extracted from BioGeoBEARS ancestral state reconstruction for the core-group. Colonization rates correspond to the proportion of transitions compared to the number of lineages existing during each 1-million year interval. The lines depicted are spline lines fitted over 100 reconstructions on the posterior distribution. The blue line is the rate of total interchanges between the Andes and Amazonia, the orange line the rate of colonization from the Central Andes toward the Northern Andes (see Supporting Information S11 for details). Dots represent the additional contribution to lineage accumulation of local diversification in three regions. Red dots represent the cumulative number of divergence events through time in Central Andes; orange dots represent the cumulative number of divergence events through time in Central Andes and Northern Andes; blue dots represent the cumulative number of divergence events through time in Central Andes, Northern Andes and Amazonia; Grey dots represent the cumulative number of divergence events through time in the entire Neotropics. B. Time-dependent speciation rates estimated on seven Amazonian clades (four in the core-group and three in the background lineages). All those clades originated during the last 8 my.

We also used the biogeographic reconstruction to estimate local diversification, namely the cumulative number of divergences inferred to have occurred exclusively in a given region. As described above, until ˜10 my ago speciation events occurring in the Central Andes fully account for the core-group diversification (no dispersal events). During the last ˜10 my, we observed a dampening of speciation events in the Central Andes (Figure 3). At the same time, following the peaks of dispersal identified above, Northern Andean and Amazonian lineages started diversifying, although the latter diversified at a slower pace than the former. This reflects the large number of dispersal events into the Northern Andes that were followed by important local diversification, for example in the genera *Hypomenitis* (17/20 species in the phylogeny occur in the Northern Andes) and *Pteronymia* (30/45 species in the phylogeny occur in the Northern Andes), or in subclades of the genera *Oleria* or *Napeogenes*. We also identified some important transitions to lowland Amazonia, for example at the origin of the *Brevioleria*-clade, during early divergence in the genus *Oleria,* or in the genus *Hypothyris* (see results on *Diversification in Amazonia*).

#### Time-stratified 328 biogeographic model

We used the results highlighted above to refine the biogeographic model by incorporating the variations of dispersal rates identified into a model accounting for time-stratified dispersal multipliers. This time-stratified model designed from rates of colonization computed above led to a significant improvement of the model (likelihoods: DECnull: −1335.802, DECstrat: −1321.805). Both ancestral state reconstructions were very congruent but the time-stratified model increased the resolution of several nodes throughout the tree (Supporting Information S11-S12). We identified one major difference in the ancestral states. From the null model the ancestral state of the subtribe Melinaeina, the first lineage to diverge, was highly unclear and the first nodes within Melinaeina were identified as Central Andean, although this was not strongly supported. Likewise, in the null model, Mechanitina, the second lineage to diverge, was inferred to have diversified in the Atlantic Forest but this was poorly supported (Supporting Information S11-S12). The time-stratified model greatly increased the resolution of all these deep nodes, inferring that both Mechanitina and Melinaeina likely initially occupied Western Amazonia. For Melinaeina and Mechanitina this result was in agreement with the BiSSE ancestral state reconstruction. The ancestral state reconstruction inferred that the Ithomiini occupied the Andes from the root, but this was very weakly supported. By contrast, all reconstructions, either BioGeoBEARS or BiSSE, inferred an Andean origin for the core-group.

## DISCUSSION

We generated one of the largest species-level phylogenies to date for a tropical insect group, the emblematic Neotropical butterfly tribe Ithomiini. With 340 out of 393 species included and a crown age of 26.4 my, this phylogeny offers a unique opportunity to investigate the dynamics of diversification of an insect group throughout the Neotropical region during the major geological and ecological events that have occurred since the Miocene. We discuss our findings below and we propose that the dynamics of multiple landscape transformations during the Miocene, and more specifically the interactions between the Andes and the Pebas system, have strongly affected the dynamics of speciation, extinction and biotic interchanges of Ithomiini butterflies in the Neotropical region.

### Early diversification at the interface of the Pebas and Central Andes: has the Pebas driven extinction?

The Ithomiini probably originated along the early Andean foothills at the transition with western Amazonia. The onset of the uplift of the eastern cordillera of the Central Andes during late Oligocene coincides with the origin of Ithomiini ^67^ and our reconstruction of the ancestral biogeographic area for the MRCA of the tribe was unable to distinguish between Central Andes or western Amazonia. The Pebas ecosystem replaced the previous western Amazonian terrestrial ecosystem from 23 to 10 my ago. Wesselingh *et al.* (^7^) described the Pebas as an ecosystem “which was permanently aquatic with minor swamps and fluvial influence, and was connected to marine environments”, and may have reached a maximum size of 1.1 million km^2^. The presence of fossil marine fishes ^68^ and molluscs ^13^ testifies to the presence of saline waters. More recently, Boonstra *et al.* ^69^ found evidence from foraminifera and dinoflagellate cysts that marine incursions reached 2000 km inland from the Caribbean sea during the early to middle Miocene during periods of high sea levels. The extent and duration of these marine influences is controversial (see ^70^ and references therein). Yet, it is undeniable that the Pebas system was not suitable for terrestrial fauna and flora, and therefore was likely to affect diversification and dispersal of the terrestrial fauna, including early Ithomiini lineages.

The timing of diversification of background lineages reveals a fast early diversification, perhaps following the colonization of South America during the pre-Pebas period – the sister clade of Ithomiini, Tellervini, is found in Australia and Papua New Guinea. Diversification was perhaps partly facilitated by an early shift to a new and diverse hostplant family, the Solanaceae, which is particularly diverse in the Neotropics ^36^, but the possible effects of new hostplants on early diversification will be difficult to distinguish from the effects of newly available landscapes. Yet, diversification rate rapidly decreased through time, driven by an increasing relative extinction rate and at a time corresponding to the replacement of the terrestrial habitats by the Pebas ecosystem, i.e., ca. 23 my ago. Although the ancestral area of Ithomiini is ambiguous, the two first diverging Ithomiini lineages (Melinaeina and Mechanitina) were clearly endemic to western Amazonia (time-stratified model in BioGeoBEARS and BiSSE reconstruction) and therefore were likely to be affected by the dramatic landscape modifications of the Miocene. However, there are uncertainties surrounding the other deep nodes and also the time when first colonization of the Andes occurred. Two scenarios can be envisioned: (1) The remaining lineages (Tithoreina, Methonina, Athesitina and the core-group) became endemic to the Central Andes (supported by the BioGeoBEARS reconstruction) and we do not know what has driven the shift of diversification at the root of the core-group; and (2) All background lineages were ancestrally western Amazonian (supported by BiSSE ancestral state reconstruction and the sister-clade Tellervini being a group restricted to lowlands) and central Andean endemicity occurred at the root of the core-group only, but extinction events in the background lineages (potentially higher in western Amazonian lineages) may have falsified the BioGeoBEARS ancestral state reconstruction. Indeed, asymmetrical extinction across different geographical regions, as suspected here, may lead to inaccurate inferences of past geographic ranges ^71,72^. In our case, if western Amazonian lineages were more prone to extinction than Andean lineages due to the presence of the Pebas, ancestral reconstruction of the distribution areas based on a phylogeny of extant taxa, i. e. those that survived extinction, will be biased towards Andean regions. Such a scenario, where background lineages were ancestrally western Amazonian, would explain the common pattern of high relative extinction rate in the background lineages and a shift of diversification process at the root of the core-group.

The idea that the Pebas may have driven extinction is well supported by a recent evaluation of an Amazonian fossil record, which pointed at a major decline of diversity in western Amazonia during the early and middle Miocene ^16^. This study concludes that mammalian diversity dropped from 11 orders, 29 families and 38 species during late Oligocene down to 1 order, 2 families and 2 species during middle Miocene (see also ^17^). These results and the pattern of extinction we found in (at least some) early Amazonian Ithomiini, which occurred during the Pebas period, strongly suggest that the late Oligocene fauna occupying western Amazonia suffered from extinction during the Pebas period. The progressive recovery of these background lineages toward the present, including positive diversification rates in some recent lineages, also concurs with the idea that the retreat of the Pebas released the constraints on diversification during the last 10 my (Supporting Information S7).

Parallel to the events occurring in western Amazonia during the Pebas period, the core-group MRCA (19.1-22.1 my old) occupied the Central Andes. This event was of major importance in shaping the diversification of Ithomiini since it is the origin of 85% of the current Ithomiini diversity. Firstly, from this event until ~10 my ago, all core-group lineages exclusively diversified in the Central Andes, meaning that from 19.1-22.1 to ~10 my ago not a single dispersal event occurred out of the Central Andes. Secondly, the core-group corresponds to a shift of diversification dynamics, characterized by a low (or zero) extinction rate and a constant speciation rate, which greatly contrasts with the slow and even negative diversification dynamics of the background lineages during the same period. Consequently, the Central Andes hosted most of the diversification during the first half of Ithomiini history. A two-fold higher Andean diversification rate was found across the whole Ithomiini, which may be mainly the consequence of the diversification rate shift found at the root of the core-group. By contrast, when considering only the core-group, Andean and non-Andean lineages had similar diversification rates. The lack of support for a general increase in diversification rate in the Andes within the core-group is also supported by analyses performed independently on different core-group subtribes. For example, in both Oleriina ^73^ and Godyridina ^19^, radiations occurred in both Andean and Amazonian genera.

### Dispersal out of the Central Andes at the demise of the Pebas

Gentry (^74^) pointed at a dichotomy observed in the geographic distribution of Neotropical plant diversity, showing that groups could be divided into Andean-centred *versus* Amazonian-centred patterns. Clades tend to be species-rich in one of these centres and relatively species-poor in the other. Antonelli & Sanmartín (^6^) coined this observation the “Gentry-pattern”. They also suggested that in the absence of a barrier between the Andes and the Amazon basin we should observe continuous interchanges between these regions. Antonelli & Sanmartín (^6^) proposed that the Pebas could be this “missing long-lasting barrier needed for creating the disjunction between Andean-centred and Amazonian-centred groups”. Therefore, in addition to the constraints on diversification discussed above, we predicted that the Pebas ecosystem should have influenced interchanges toward or across western Amazonia.

Our results conform surprisingly well to the scenario proposed by Antonelli & Sanmartín (^6^). Ithomiini are Andean-centered with more than a half of their current diversity occurring in the Andes (see also ^75^). Here we show that interchanges have been virtually absent during the Pebas period, with a period as long as 9-12 my without interchanges. However, rates of interchanges from the Central Andes toward the Northern Andes and Amazonia suddenly peaked ~10 my ago (between 13-8 my ago) and more recently (4-0 my ago). The Western Andean Portal (WAP) is a low-altitude gap that separated the Central Andes and the Northern Andes until 13-11 my ago, and which may have connected the Pebas system and the Pacific Ocean (^6^). The closure of the WAP may have allowed multiple colonizations of the Northern Andes facilitated by the presence of connecting higher altitude habitats. In parallel, between 10-8 my ago the Pebas system was drained eastward, leading to the formation of the present-day configuration of Amazonian drainage basin. It was accompanied by the expansion of terrestrial forest habitats in western Amazonia. This corresponds precisely to the timing at which core-group lineages colonized western Amazonia and then diversified.

### Diversification across the whole Neotropics following the demise of the Pebas

We found a strong dampening of local speciation in the Central Andes during the last 10 my. However, colonizations following the retreat of the Pebas system were followed by large local bursts of diversification within the Northern Andes and Amazonia. As an illustration, from our biogeographic reconstruction, 69 divergence events occurred strictly in the Central Andes in the core-group from 20 my ago until present-day. However, multiple independent dispersal events followed by local diversification lead to the exact same number of divergences occurring strictly in the Northern Andes during the last 9 my only. The genera *Hypomenitis* and *Pteronymia,* for example, diversified extensively within the Northern Andes with 23 and 53 valid species respectively. We also identified four Amazonian radiations in the core-group and three in the background lineages. Lineages that dispersed into the Northern Andes and Amazonia after the demise of the Pebas system probably benefited from a large range of free ecological niches, including a diversity of host-plants that had already diversified or that radiated concomitantly. Two of the background Amazonian radiations, the genus *Melinaea* and the clade *Mechanitis* + *Forbestra,* showed increasing speciation rate toward the present, due in the former case to the shift detected by MEDUSA (*Melinaea*-group). The recent and dramatic increase in diversification rate of the *Melinaea*-group, which produced at least eight species and 50 subspecies ^43^ in just 1 my, is particularly intriguing. This radiation may be interpreted as support for an effect of recent climatic fluctuations during the Pleistocene on the diversification of this group (as well as the *Mechanitis* + *Forbestra* clade), although ecological drivers of speciation classically invoked in mimetic butterfly diversification, such as colour pattern and hostplant shifts, cannot be ruled out ^42,43^. Five other Amazonian radiations showed diversification rates decreasing through time, meaning that diversification was highest just after the retreat of the Pebas. Recent radiations in western Amazonia that post-date the Pebas period have been repeatedly reported. For example, in the genus *Astrocaryum* (Arecaceae), the upper Amazonian clade started diversifying only ~6 my ago ^76^. In *Taygetis* butterflies, Amazonian lineages show rapid diversification during the last 7-8 my ^35^. Such convergent timing of diversification in western Amazonia strongly supports the scenario of a post-Pebas recovery of terrestrial habitats, which triggered dispersal followed by local diversification.

### Conclusion

Our research shows that the timing of diversification and biogeographic interchanges in Ithomiini butterflies are tightly associated with the turnover of ecological conditions that occurred during the Miocene. Our findings suggest that the ecological turnover that first accompanied the expansion of the Pebas system has led to a decline of diversification, potentially driven by increasing extinction, in early lineages adapted to the ecological conditions that existed during the Oligocene in western Amazonia. Such a decline of diversity has also been documented in the fossil record ^16^, which calls for further investigations on the role of the Pebas in driving extinction during the Miocene. By contrast, lineages that colonized the Central Andes 20 my ago rapidly diversified. However, during the entire existence of the Pebas, these lineages remained trapped in the Central Andes (at least 9-12 my without dispersal events out of the Andes). The closure of the West Andean Portal, connecting the Central and North Andes, and the associated demise of the Pebas (10-8 my ago), apparently released these long-lasting barriers, allowing interchanges with the Northern Andes and Amazonia and opening new opportunities for diversification. As a result of these multiple events, major differences appear between the different faunas. Central Andean lineages started diversifying early (at least 19 my ago), allowing species to accumulate over a long period of time, but diversification slowed down during the last 10 my. In contrast, the Northern Andean fauna is recent (13-11 my old at most), driven by multiple colonization events sometimes followed by important bursts of diversification. In parallel, some Amazonian lineages may be old (late Oligocene), but modern diversity almost entirely arose during the last 8-10 my, after the demise of the Pebas ecosystem. Taken together, all this information points to a robust scenario for Neotropical diversification, which highlights the role of Miocene ecosystem turnover in determining the timing of interchanges, speciation and extinction in the world’s most biologically diverse region.

## MATERIAL AND METHODS

### Time-calibrated phylogeny

#### Molecular data

We compiled sequences of 1460 Ithomiini individuals (Supporting Information S1) that included sequences newly generated for this study and previously published sequences ^19,21,26,27,32,47,52,77^. We used a concatenation of nine gene fragments, a mitochondrial fragment spanning genes COI-tRNA-COII, and fragments of nuclear genes EFIα, Tektin, CAD, RPS2, MDH, GAPDH, representing a total of 7083 bp ^78^. Primers and PCR conditions followed ^32 and 78^. We obtained at least one gene fragment for 340 species out of 393 currently known in the group, which represents 87% of the known species richness of the tribe. For each species we produced the consensus sequence of all sequences belonging to individuals of that species to obtain the longest sequence possible. We added 41 outgroups, which spanned all Danainae genera as well as representatives of the main Nymphalidae clades. In total, seven concatenated genes from 381 taxa were used to generate the time-calibrated phylogeny of the Ithomiini.

#### Tree topology and time calibrations

First, we generated a phylogeny under maximum likelihood inference (ML), using IQ-Tree software as implemented in the W-ID-TREE server ^79 80^ in order to obtain a tree topology (Supporting Information S2). Then, using BEAST v1.8.2 ^81^, we time calibrated this tree by enforcing the ML topology and preventing BEAST from searching for a new topology in the xml file and following the calibration procedure described below.

Gene partitioning and substitution models were estimated using PartitionFinder v.1.1 ^82^. We divided all gene fragments into codon positions and allowed all partitions to be tested. Only substitution models available in BEAST were tested. The models of linked partitions had a better fit than unlinked partition schemes, hence the former was used. The best linked partition scheme contained 13 partitions (Supporting Information S3).

Branch lengths were estimated using BEAST v.1.8.2 ^81^ with a birth-death tree prior and a lognormal relaxed molecular clock for each gene partition. In order to time-calibrate the tree we also used a combination of secondary calibrations and host-plant calibrations (Supporting Information S4). Four secondary calibration points were retrieved from Wahlberg *et al.* (^57^)’s phylogeny of Nymphalidae genera and were placed only outside of Ithomiini. We used uniform distribution priors, corresponding to the 95% HSPD inferred by Wahlberg *et al.* (^57^). Host-plant calibrations were used as constraints only within the Ithomiini. Almost all Ithomiini species feed on Solanaceae with a relatively high specificity. A phylogeny of the Solanaceae was published by Särkinen *et al.* (^83^), and recalibrated by De-Silva *et al.* (^48^). We identified six relevant hostplant clades (Supporting Information S4), which were used as maximum age constraints. Priors for host-plant calibration followed a uniform distribution. The minimum value of the uniform was 0 (present). The maximum value was the upper boundary of the 95% HSPD of the stem age of the host-plant subclade on which the calibrated Ithomiini lineage feeds. To get a starting tree suitable for time-calibration priors, the ML tree was ultrametrised and rescaled using PATHD8 ^84^ and Mesquite v.3.2 ^85^.

We performed two independent runs of BEAST v.1.8.2 ^81^ on the CIPRES server ^86^ of 87 million generations, sampling every 10000. Using Tracer v.1.6 we checked that the two runs had converged and the parameter’s ESS values. Both runs were combined using Logcombiner v.1.8.3 ^81^, applying a 15% burn-in for each run. Finally using TreeAnnotator v.1.8.3 ^81^ we extracted the Maximum Clade Credibility tree with median branch lengths and we computed the posterior probabilities of each node (Supporting Information S5). The outgroups were pruned and the remaining tree was used for the following analyses (hereafter, MCC tree).

### Diversification rates

#### Time

We investigated the dynamics of speciation and extinction rates through time and across the phylogeny. We chose a combination of three methods to infer the dynamics of diversification: MEDUSA ^59^ followed by time-dependent models of diversification proposed by Morlon *et al.* (^60^) and Stadler (^61^, TreePar). MEDUSA is a maximum likelihood method that uses stepwise AIC to automatically identify the number and the position of different diversification processes that maximize the fit of the model to the tree. However, MEDUSA makes the strong assumption that rates of speciation and extinction are constant through time. To relax this assumption, we combined MEDUSA with two alternative methods. (1) Morlon *et al.* (^60^) developed a method that allows both speciation and extinction rates to vary as a function of time and across lineages, and where extinction is allowed to be higher than speciation, a situation leading to declining diversity. However, it does not automatically detect rate shift points, which therefore have to be specified by the user. (2) TreePar ^61^ accommodates models where diversification rates can vary at points in time but are constant between these points. Similarly to Morlon *et al.* (^60^)’s method, TreePar does not automatically detect shifts within a tree. We proceeded as follows: we ran MEDUSA on the whole tree (MCC tree) in order to partition the tree into different diversification processes but rate estimates were ignored. Instead, we used the partition inferred by MEDUSA to estimate diversification rate (speciation - extinction) and turnover rate (extinction/speciation) by fitting time-dependent models of diversification using Morlon *et al.* (^60^)’s method and TreePar ^61^. MEDUSA detected two shifts from the background diversification rates (see Results and Supporting Information S6-7): 1) one shift at the root of a large clade hereafter referred to as the core-group, and 2) one shift at the root of a subclade of the genus *Melinaea* that, for simplicity, is referred to as the *Melinaea*-group. We fitted time-dependent models using the method provided in Morlon *et al.* (2011) on all four possible partitions: no shift, one shift (either the coregroup or the *Melinaea*-group) and both shifts. Sampling fraction was indicated for each partition. Comparing the fit of all partitions allowed us to confirm that the two shifts identified by MEDUSA were indeed significant. In each case the stem branch of the shifting clade was included in the subclade, as designed in the method ^60^, but we excluded the stem branch of the background tree. For each distinct part of the tree (background tree, *Melinaea*-group and core-group) we fitted the following models: constant speciation (no extinction), time-dependent speciation (no extinction), constant speciation and extinction, time-dependent speciation and constant extinction, constant speciation and time-dependent extinction, time-dependent speciation and extinction. In the cases of time-dependent rates we fitted an exponential dependence to time. Sampling fraction was specified for each of the three partitions. All models were compared using AIC scores. The model with the lowest AIC score that was significantly different from the null model of constant speciation rate was used to plot the diversification rate through time. Finally we ran TreePar ^61^ in order to obtain a second, independent, estimation of diversification and turnover rates across time. TreePar uses a vector of speciation times, which allowed us to run it separately on the core-group and the background tree. For the background tree we added the times of divergence at which the core-group and the *Melinaea-group* diverged from the background to keep track of these cladogenetic events. We split time into time bins of five million years, for which diversification and turnover rates were estimated. We used these estimates to obtain a second estimate of diversification rate “through time” that could be compared with Morlon *et al.* (^60^)’s method. We did not fit TreePar on the *Melinaea*-group since it is only 1 million year old, and therefore does not even span an entire time bin. We allowed diversification rate to be negative but we did not allow mass-extinction events.

In addition to MEDUSA, Morlon *et al.* (2011) and TreePar we also performed analyses with BAMM v 2.5.0 ^62^. We ran BAMM for 30 million generations sampling every 3000 generations, using genus level sampling fraction. The results of the MCMC were analyzed in the R package BAMMTOOLS ^87^. First we checked for MCMC convergence using trace plots and ESS statistics using the R package CODA, after removing 10% of the chain as burnin. Second we investigated the number and configurations of diversification rate shifts (Details and results can be found in Supporting Information S8).

#### Diversification in the Andes

To investigate the pattern of diversification in the Andes with respect to rest of the Neotropical regions, we classified species as Andean or non-Andean, based on a combination of published^19,21,27,48,52,68^ and unpublished (i.e., databases generated from museum collections and our own field collection) georeferenced distribution data and elevation ranges of species. In general species can be unambiguously classified to either of the two categories, because Andean species are never found in the lowlands, whereas species that occur in the lowlands and that sometimes also occur at the Andean foothills never occur above ca. 800m, and are therefore classified as non-Andean. Species that do not occur in the Andean region (e.g., Atlantic Forest) are obviously considered as non-Andean. Species biogeographic distribution was used as character states. Then we used trait-dependent models of diversification ^19,88^ to compare the rates of speciation, of extinction and of transition between the Andean area and the non-Andean regions. We used the ClaSSE model ^63^, which accounts for up to 10 parameters (2 speciation rates without character state change, 4 cladogenetic transition rates, 2 extinction rates and 2 anagenetic transition rates). However, we constrained parameters that were not biogeographically meaningful to zero ^19^. Those include the anagenetic transition rates, considering that transition rates from one region to the other were always accompanied by a speciation event, and the cladogenetic transition rates involving a transition in both descendant lineages since we considered the scenario as unrealistic. We therefore ended up with at most 6 parameters. We tested all models with only one parameter (speciation, extinction, transitions) free to vary as well as all models combining two or more parameters free to vary. This allows comparing the relative contribution of one biogeographic model to the others, as well as their combination since they are not mutually exclusive. Models were compared using AIC scores. All models were fitted on the MCC tree.

Furthermore, we considered two important sources of potential biases. (1) The major shift of diversification at the root of the core-group could affect our results of trait-dependent diversification models. Thus we fitted ClaSSE on both the whole tree and the core-group alone to compare the results. (2) We performed additional analyses to test the hypothesis that a hidden character not considered here explained the pattern of trait-dependent diversification measured with ClaSSE. To do so we fitted the model HiSSE ^64^ HISSE results were fully congruent with the model ClaSSE and our interpretation. The details about the models fitted, results and interpretations can be found in Supporting Information S9.

Finally, we also conducted ancestral state reconstructions based on the models of trait-dependent diversification. Since ancestral state reconstruction is not available for ClaSSE models we used the BiSSE model ^65^ for this purpose. BiSSE also includes 2 speciation rates and two extinction rates, but it allows the transitions to occur only along branches (anagenetic transition rates). Therefore, we fitted the BiSSE model corresponding to the best fitting ClaSSE model, and we used these parameters to infer the ancestral states at the nodes of the phylogeny. This ancestral state reconstruction was compared to that obtained using the historical biogeography analyses outlined below. All these analyses were performed on both the whole phylogeny and the core-group only, to account for the diversification rate shift identified by our time-dependent diversification analyses.

#### Diversification in Amazonia

We further investigated the pattern of diversification in Amazonia during the post-Pebas period. The Amazonian basin appears to be the second most important place for diversification after the Andes and there is a longstanding hypothesis that speciation in this region has been driven by climatic fluctuations during the Quaternary ^9^. An interpretation of this scenario is that speciation rate should increase during the last 2.5 million years ^12^. To test this hypothesis we identified the major Amazonian diversification events, i.e. clades which nodes were inferred to be almost all Amazonian from the BioGeoBEARS ancestral state reconstruction. We fitted a model of time-dependent speciation rate (no extinction) to see whether speciation rates increased through time (supporting a recent diversification potentially caused by Pleistocene climatic fluctuations) or decreased through time (supporting radiations accompanying the post-Pebas recolonizations).

### Historical biogeography

We proceeded in three steps to reconstruct the historical biogeography of Ithomiini. First we performed an ancestral state reconstruction using a model with refined area adjacency but uniform dispersal multipliers (null-model). Second, we used the results of this model to compute rates of dispersal between specific regions per million years. This allowed us to test some biogeographic hypotheses but also to identify relevant time frames for which dispersal probabilities might vary. Third, we implemented a time-stratified model designed from the previous information to refine our biogeographic reconstruction.

#### Biogeographic null model

We inferred the historical biogeography of Ithomiini using BioGeoBEARS v.0.2.1 ^66^. We divided the Neotropics into nine distinct biogeographic regions (Supporting Information S11): 1) Central America, 2) Caribbean Islands, 3) lowlands on the western part of the Andes, including the Magdalena valley, 4) Northern Andes that comprise the western and eastern Ecuadorian and Colombian cordilleras and the Venezuelan cordillera, 5) Central Andes, 6) western Amazonia, 7) eastern Amazonia, 8) Guiana Shield and 9) Atlantic Forest (Supporting Information S11). In this model, we constrained the combinations of areas to avoid unrealistic distributions (e.g., disjunct distributions) but all dispersal multipliers were set to 1 to avoid biasing the ancestral state reconstruction. We compared the models DIVALIKE and DEC as implemented in BioGeoBEARS v.0.2.1 ^66^ using log likelihoods. We used the best fitting model (DEC) for the following analyses.

#### Biogeographic interchanges within the coregroup

One important hypothesis is that the Pebas influenced the interchanges among biogeographic regions, especially toward or across western Amazonia. To test this hypothesis, we computed “rates” of colonization from BioGeoBEARS ancestral-state reconstructions performed with the null-model. For 100 trees randomly sampled from the BEAST posterior distribution we applied the null-model with BioGeoBEARS and reconstructed the ancestral states. For each tree, we extracted the state with the highest probability at each node. When a descendent node had a range different from the ancestral node, we considered the middle of the branch connecting these nodes as the timing of the event and recoded it. Then for each million-year interval, we computed the proportion of specific state transitions compared to the maximum number of lineages existing during this interval. Hence, for each million-year interval we obtained a proportion of lineages that dispersed for example from the Central-Andes toward the Northern-Andes or from the Andes toward the Amazonia, which we refer to as “rates” of colonization. Therefore, for each million-year interval and each state transition we obtained a distribution of values. We fitted a spline line on this distribution against time to assess the pattern of variation of colonization rates through time. That way we computed rates of colonization between Andean and non-Andean regions, between the Andes and the Amazonia, between the Central and the Northern Andes, between the Atlantic forest and the other regions, and between Central America and the rest of the Neotropics. These rates were computed only on the core-group, because (1) this group contains 85% of Ithomiini species, (2) the diversification process is homogeneous throughout the clade and (3) we found high extinction in background lineages, which may falsify the ancestral state reconstruction in these lineages, which were therefore excluded. We focused more specifically on the timing of transitions between the Andes and the Amazonian basin and the timing of transitions from the Central Andes (ancestral Andean region of the core-group) to the Northern Andes, because a strong influence of the Pebas and the WAP should have affected these rates of interchanges (see Supporting Information S11 for the complete results).

Additionally, we recorded the divergence times of nodes inferred to be strictly in either the Central-Andes, the Northern Andes or Amazonia (the region comprising the upper and lower Amazon and the Guiana shield). We excluded all nodes inferred to be in more than one of these areas (for Amazonia, only all combinations of the three areas above were considered) so that we obtained times of divergences inferred to have occurred strictly within the region considered (which we call local diversification). These divergence times were obtained from the unconstrained BioGeoBEARS reconstruction (DEC) on the MCC tree and we used them to represent lineage accumulation through time (exclusively due to speciation) in each region.

#### Time-stratified biogeographic model

We used our estimations of colonization rates among the major biogeographic regions to design a time-stratified biogeographic model and improve the resolution of our biogeographic reconstruction (Supporting Information S11). We created four time frames: i) 0-4, ii) 4-8, iii) 8-13, and iv) 13-30 my ago, where dispersal multipliers between areas reflected the colonization rate variations identified previously. This ancestral state reconstruction was compared to the null model as well as to the ancestral state reconstruction obtained from trait-dependent diversification models (Supporting Information S10).

## Acknowledgements

This project was funded by an ATIP (CNRS, France) grant awarded to ME, with LDS as a postdoc. NC was funded by a doctoral fellowship from Ecole Doctorale 227 (France). ME acknowledges additional funding by the ANR grant SPECREP (ANR-14-CE02-0011-01). We thank the authorities of Peru, Ecuador and Brazil (SISBIO n° 10438-1) for providing research and collection permits, as well as many assistants for their help in the field. Molecular work was performed at the GenePool (University of Edinburgh, UK), CBMEG-Unicamp (Brazil) and the Service de Systématique Moléculaire UMS2700 of the MNHN (France). We are grateful to Niklas Wahlberg for providing unpublished sequences of *Greta diaphanus.* We thank Fabien Condamine and Hélène Morlon for constructive discussions about time-dependent diversification analyses. AVLF thanks CNPq (grant 303834/2015-3), RedeLep-SISBIOTA-Brasil/CNPq (563332/2010-7), National Science Foundation (DEB-1256742), FAPESP (grants 2011/50225-3 and 2012/50260-6) and USAID (Mapping and Conserving Butterfly Biodiversity in the Brazilian Amazon).

## Author contributions

NC and ME conceived the study, with contribution from KRW, GL and AVLF. All co-authors provided specimens and sequences. NC, ME, FPP, CFA, DLDS performed the labwork. NC performed the analyses. NC wrote the paper with major contributions from ME, and contributions from all co-authors.

## List of supporting information

S1. List of all individuals and species used in this study, and biogeographic distribution of species.

S2. Gene fragment partitions and substitution models associated obtained by Partition Finder v.1.1

S3. Node constraints used for time-calibration of the tree.

S4. Maximum likelihood tree obtained with IQ-tree, with bootstrap support indicated at the nodes.

S5. Bayesian time-calibrated tree obtained with BEAST v.1.8 with median node ages and 95%HPD indicated at the nodes.

S6. Full results of diversification analyses.

S7. Testing the effect of diversification rate heterogeneity within the background on diversification rate estimates

S8. Diversification analysis with BAMM.

S9. Trait-dependent diversification using HiSSE.

S9. Ancestral state reconstruction performed using trait-dependent diversification models

S10. Rates of colonization between different region, computed on the core-group and used to design a time-stratified biogeographic model.

S11. Results of biogeographic ancestral state reconstruction obtained with BioGeoBEARS and using the “null” or time-stratified model.

